# Differential effects of ankle constraints on foot placement control between normal and split belt treadmills

**DOI:** 10.1101/2022.07.18.500502

**Authors:** Mayra Hos, Lieke van Iersel, Moira van Leeuwen, Sjoerd M. Bruijn

## Abstract

Mediolateral ankle moment control contributes to gait stability. Ankle moments can be constrained by walking with a shoe with a ridge underneath the sole, narrowing the mediolateral support surface. In our previous study, such ankle moment constraints resulted in an increased step width and a decrease in the degree of foot placement control, as defined by the percentage of variance in foot placement that can be explained by CoM state. However, since our previous study was performed on a split-belt treadmill and the narrow ridge could fit inside the gap between the belts, it is not evident whether these effects can be attributed to the constrained ankle moment control or to avoidance of this gap. Therefore, we investigated if the effects of ankle moment constraints are dependent on whether participants walk on a normal treadmill or a split-belt treadmill. We included fourteen healthy young adults. Walking with constrained ankle moment control resulted in a wider step width on both treadmills. Yet, the increase in step width was larger on the split-belt treadmill compared to on the normal treadmill. We only found a decreased degree of foot placement control on the split-belt treadmill, whilst the degree of foot placement control increased on the normal treadmill. We conclude that the effects of ankle moment constraints reported in our previous study were confounded by the use of a split-belt treadmill. For future research, we recommend using a normal treadmill whenever possible, because the gap in a split-belt treadmill might affect gait parameters.

## 1. Introduction

In bipedal locomotion, the Centre of Mass (CoM) is relatively high above a relatively small and continuously changing Base of Support (BoS). This makes stability control complex, as a perturbation can rapidly result in instability (Bruijn & Van Dieën, 2018; Reimann et al., 2018). The vertical projection of the CoM remains medial from the BoS during single-legged support. To prevent falling, the legs must be placed accurately to accommodate changes in the CoM kinematic state (Reimann et al., 2018; Winter et al., 1990).

The changing of the base of support via foot placement has been coined the foot placement strategy (Bruijn & Van Dieën, 2018; Hof et al., 2007; Reimann et al., 2018). When increasing step width, larger changes of CoM movement can be accommodated (Lencioni et al., 2020; Reimann et al., 2018). Moreover, maintaining a high degree of step-by-step foot placement control with respect to variations in CoM kinematic state ensures stability, also at narrower step widths (Mahaki et al., 2019; Perry & Srinivasan, 2017). In addition to the foot placement strategy, individuals can control their ankle moment so as to shift the center of pressure (CoP) under the foot during the stance phase. Through the displacement of the CoP, the ground reaction force (GRF) provides a more beneficial torque with respect to the CoM, compensating for “errors” in foot placement (Bruijn & Van Dieën, 2018; Hof et al., 2007; Reimann et al., 2018; van Leeuwen et al., 2021, 2022).

In recent work, we (Van Leeuwen et al., 2020) investigated the effects of constraining ankle moment control by a shoe (LesSchuh) with a narrow, yet flexible, ridge underneath the shoes’ soles (somewhat like skates). Constraining ankle moment control with LesSchuh resulted in an increased step width and a decreased degree of foot placement control (as defined by the percentage of foot placement that can be explained from the CoM state), which is the relative explained variance of foot placement control (Wang & Srinivasan, 2014). However, along with the constrained ankle moment control, these results might have been affected by fear of stepping in the gap between the belts of the split-belt treadmill the study was performed on.

In the current study, we investigated whether the effects we reported earlier were confounded by the use of a split-belt treadmill. Therefore, we compared conditions on a normal treadmill and on a split-belt treadmill. In our previous study, constraining ankle moment control while wearing LesSchuh resulted in an increased step width and a decrease in the degree of foot placement control on a split-belt treadmill (Van Leeuwen et al., 2020). Despite a potential fear of stepping into the gap, we consider these findings to reflect general adaptations to constrained ankle moment control. The diminished degree of foot placement control may be explained by a steering role of ankle moment control of subsequent foot placement. Wider steps could be adopted to maintain stability despite the constraints on ankle moment control. Therefore, despite the absence of a gap while walking on a normal treadmill, we hypothesized that constraining ankle moment control would result in an increased step width and a diminished degree of foot placement control, both when walking on a normal and on a split-belt treadmill.

## 2. Methods

### 2.1. participants

Fifteen young adults, without injuries or other constraints which might affect gait, participated in this study. Data of one participant were not included in the statistical analyses, due to measurement errors. The remaining participants (3 men, 11 women, age 22.4 (2.9) years (mean(sd)), weight 67.3 (12) kg, length 1.72 (0.08) m) are included in the statistical analysis. All participants signed informed consent, and the protocol was approved by the local ethical review board (VCWE-2019-108).

### 2.2. Research design

Participants walked on two different treadmills (Splitbelt, C-Mill; Motek-Forcelink, Amsterdam, Netherland) with two types of shoes (regular, LesSchuh), making for a total of four conditions (see table 1). The Splitbelt is a treadmill with a small gap between the two belts. The C-mill is a regular treadmill. The Regular shoe is a sneaker with a flat sole. LesSchuh is the same kind of sneaker, however, a narrow ridge (one centimeter) is attached to the sole, see Figure 1 (van Leeuwen et al., 2020, p. 5). This ridge constrains ankle moment control during walking.

**Table 1.**
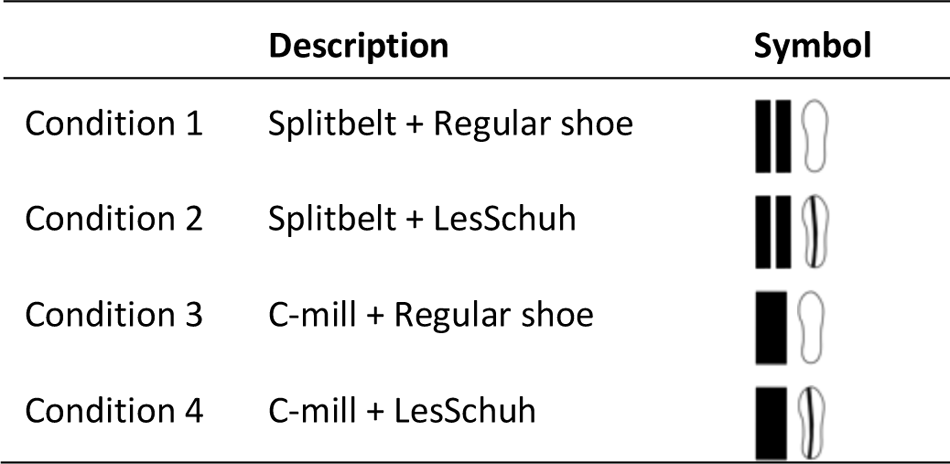
Conditions.

|  | Description | Symbol |
| --- | --- | --- |
| Condition 1 | Splitbelt + Regular shoe |  |
| Condition 2 | Splitbelt + LesSchuh |  |
| Condition 3 | C-mill + Regular shoe |  |
| Condition 4 | C-mill + LesSchuh |  |

**Figure 1.**
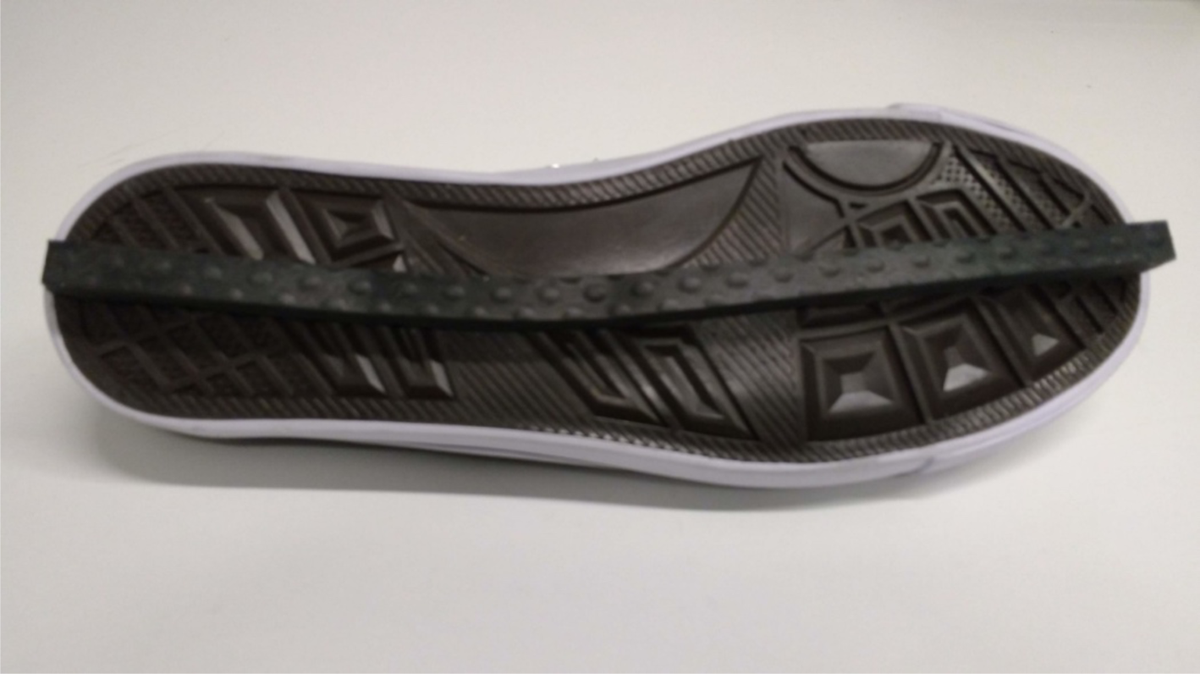
LesSchuh. The narrow ridge has a width of one centimeter and bends along with the sole, allowing for normal anterior-posterior roll-off, yet constraining mediolateral center-of-pressure modulation. Reprinted from “Active foot placement control ensures stable gait: Effect of constraints on foot placement and ankle moments”, by A.M. van Leeuwen, J.H. van Dieën, A. Daffertshofer and S.M. Bruijn (2020), *PLOS ONE, 15* (12), p. 5.

### 2.3. Procedure

First, the preferred step frequency was determined while walking at a speed of 1.25 × √(leg length [m]) m/s (Hof, 1996). During all subsequent trials, subjects walked at this step frequency, as dictated by a metronome.

For both treadmills we used the same set-up (see Figure 2), consisting of the treadmill and two Optotrak camera arrays (Northern Digital Inc, Waterloo Ontario, Canada), which recorded kinematics of three cluster markers attached to the thorax and the back of both shoes (sampling rate 50 Hz). Before the measurements, anatomical landmarks were defined with a pointer. We have used these landmarks to determine a proxy for the CoM (processus xiphoideus, incisura jugularis, processus spinosus Th7 and C7), the orientation of the feet (phalanx distalis II, tuber calcanei, malleolus medialis and malleolus lateralis) and step width (tuber calcanei).

**Figure 2.**
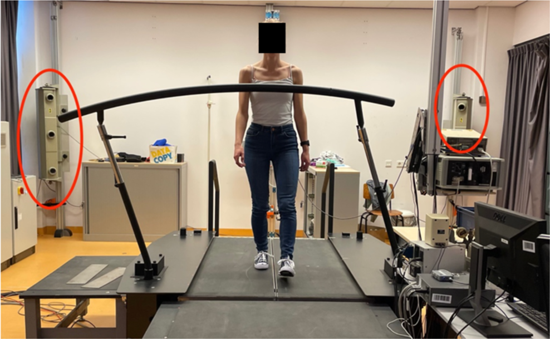
Experimental Set-up Splitbelt. The red circles in this figure point out the Optotrak camera arrays.

Each condition lasted twelve minutes, of which the first two minutes were regarded as warming-up. We used counterbalanced randomization for the conditions. On both treadmills participants started on the same shoes. During the measurement period, we informed the participant of the remaining walking time at five and nine minutes. Between every condition, a rest period was included. During this period, participants were asked to change their shoes.

### 2.4. Data processing

Using the heel markers’ vertical position and velocity, we determined the moments of heel strike and toe-off (Pijnappels et al., 2001). Midstance was defined as the midpoint in time between heel strike and toe-off. At midstance, we calculated step width as the distance between the calcanei of the two feet.

The degree of foot placement control was calculated as the relative explained variance of a regression model which relates mediolateral foot placement to mediolateral CoM position and velocity, in accordance with (Bruijn & van Leeuwen, 2020; van Leeuwen et al., 2020; Wang & Srinivasan, 2014). The data and code for analysis is available on Zenodo https://doi.org/10.5281/zenodo.7188711.

As secondary outcome measures, we determined stride frequency by calculating the mean stride frequency over the measurement period and the toe-out angle at midstance.

### 2.5. Statistics

A power analysis (a = .05; statistical power = .80) indicated that a minimum of nine participants were required for this study (effect size for step width in data of (van Leeuwen et al., 2020a)= 0.83).

We conducted repeated measures ANOVAs with factors Shoe [Regular, LesSchuh] and Treadmill [Splitbelt, C-mill] for step width and foot placement control. All statistical analyses were performed with Jamovi version 1.6.23.0 (The jamovi project, 2021), and p<0.05 was considered significant.

## 3. Results

For step width, we found significant effects of Treadmill and Shoe, as well as their interaction (see Table 2). The interaction showed a greater effect of LesSchuh on the split-belt treadmill, compared to on the C-mill (see Figure 3).

**Table 2.** Results of statistical tests. Significant effects are displayed in **bold**.

|  |  | Treadmill | Shoe | Treadmill*Shoe |
| --- | --- | --- | --- | --- |
| Step width | F(1,13) | 108.9 | 27.6 | 15.3 |
|  | p | **< 0.001** | **< 0.001** | **0.002** |
| $R^2$ | F(1,13) | 49.969 | 0.492 | 0.496 |
|  | p | **< 0.001** | 0.496 | **< 0.001** |

**Figure 3.**
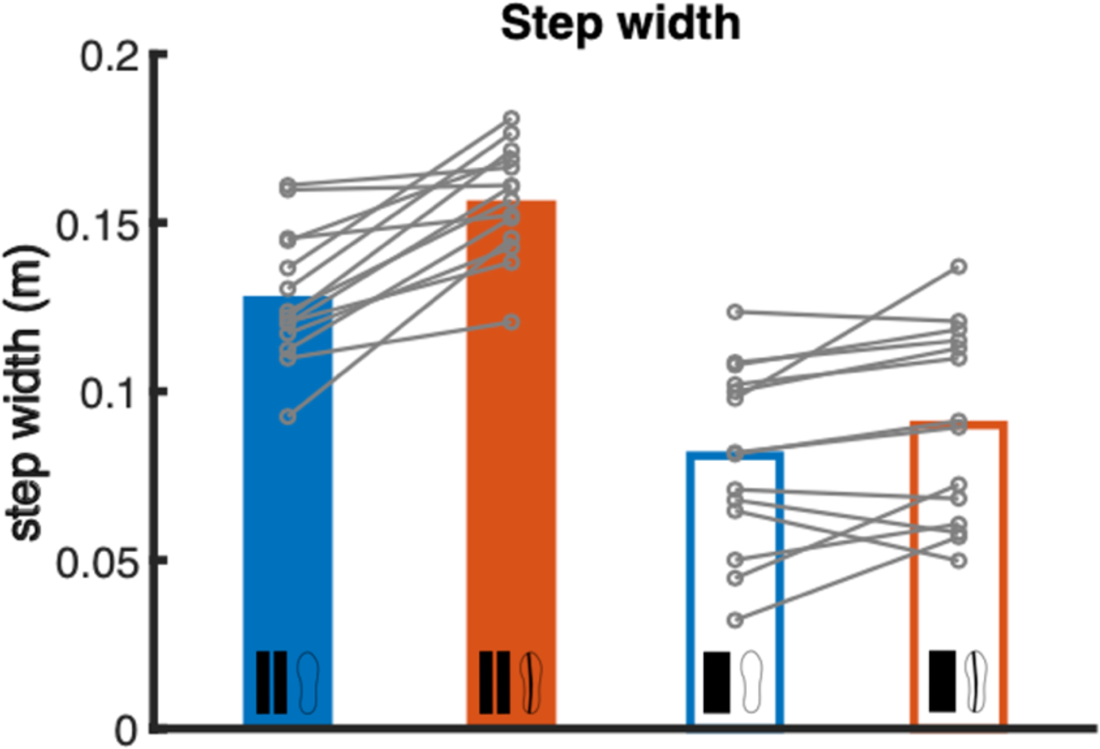
Effects of Treadmill and Shoe on step width. The grey dots and lines represent the individual data, bars represent averages.

For R^2^, we found a significant main effect for Treadmill and a significant interaction effect (see table 2). The interaction shows opposite effects of ankle moment constraint for the split-belt treadmill and the C-Mill. On the split-belt treadmill, LesSchuh decreased the degree of foot placement control compared to wearing regular shoes (p<0.05), whereas on the C-mill the degree of foot placement control increased when walking with LesSchuh (p<0.01) (see Figure 4).

**Figure 4.**
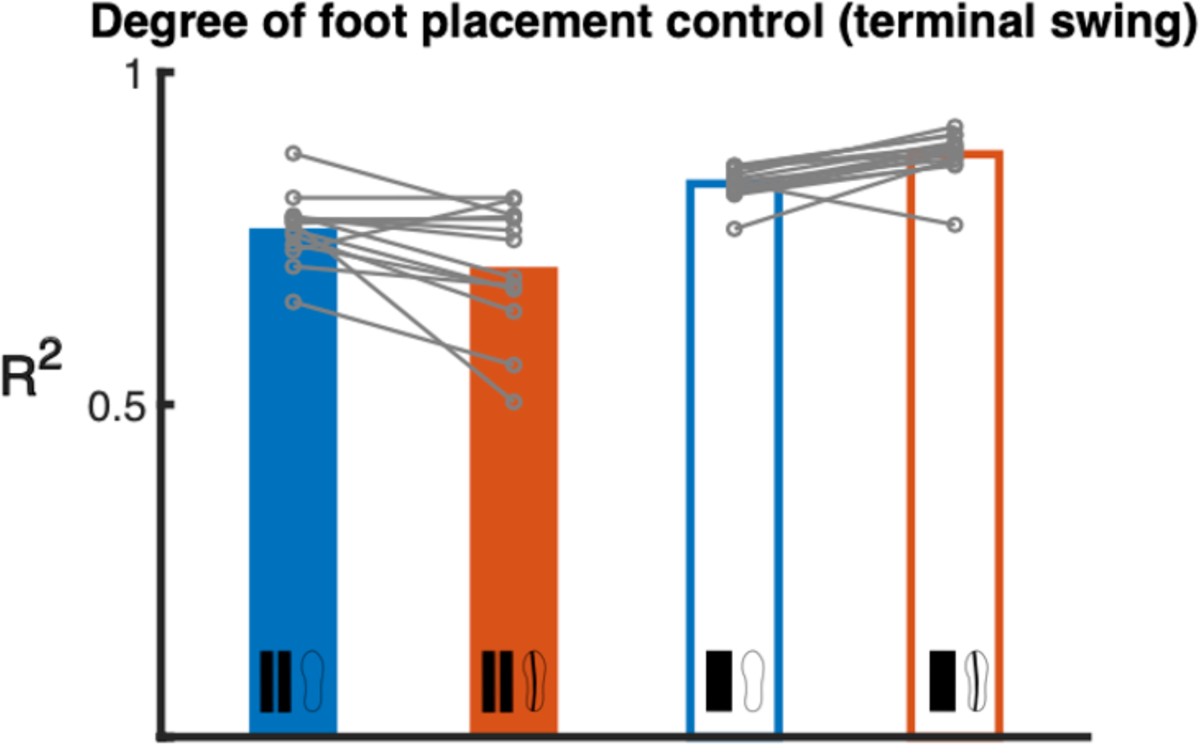
Effects of Treadmill and Shoe on R^2^. The grey dots and lines represent the individual data, bars represent averages.

For the secondary outcome measure stride frequency and toe-out angle, we found significant main effects for Treadmill and Shoe (see table 3, Figure 5 and Figure 6).

**Table 3.**
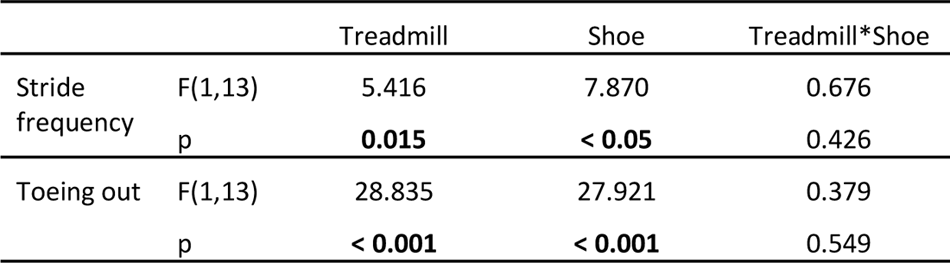
Results of statistical tests. Significant effects are displayed in **bold**.

**Figure 5.**
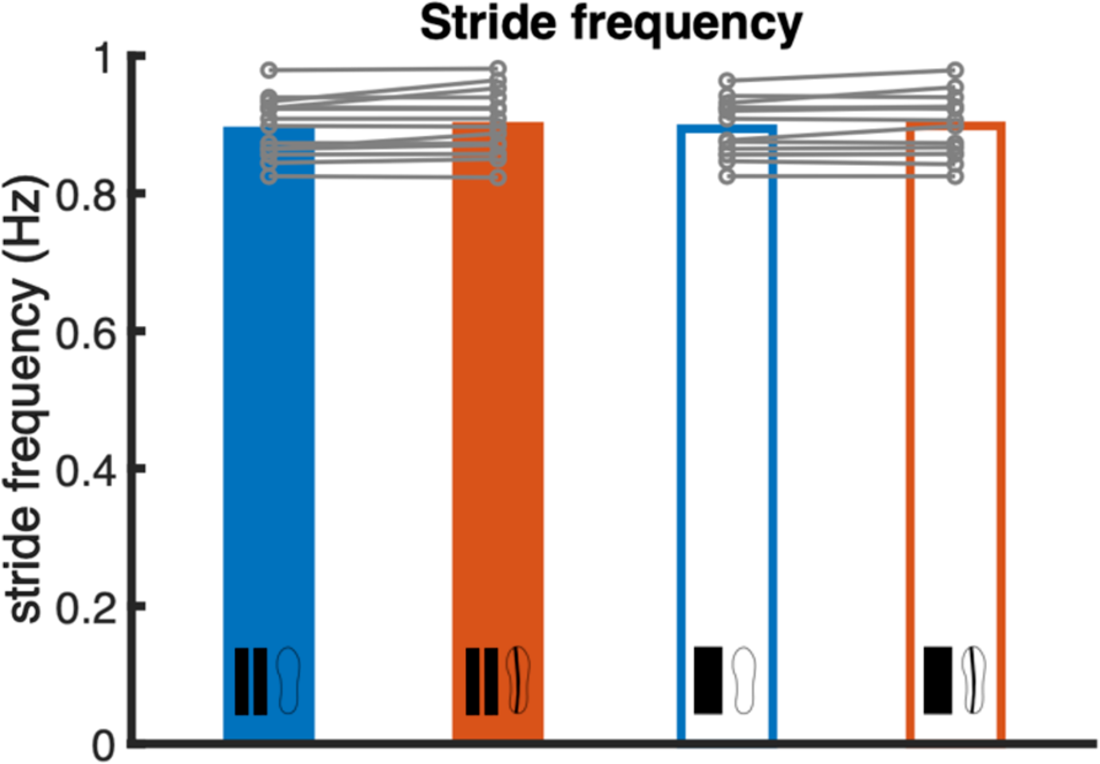
Stride frequency. The grey dots and lines represent the individual data, bars represent the average.

**Figure 6.**
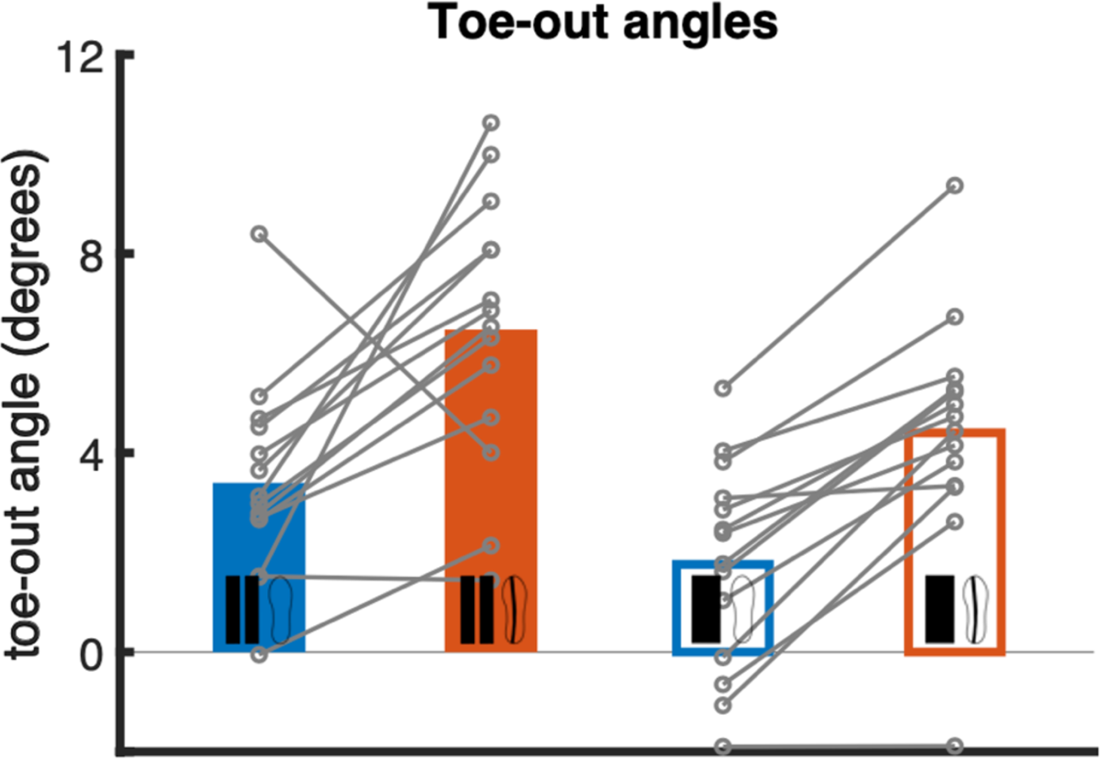
Toeing out. The grey dots and lines represent the individual data, bars represent the average.

## 4. Discussion

In a recent study, we concluded that constrained ankle moment control resulted in a greater step width and a decreased degree of foot placement control (Van Leeuwen et al., 2020). However, that study was performed on a split-belt treadmill, and thus, the observed effect might be partially explained by the gap between the belts. To investigate whether constraining ankle moment control affects step width and foot placement control, regardless of treadmill type, we conducted our measurements on a split-belt treadmill and a C-mill (normal treadmill).

We hypothesized that constraining ankle moments, by wearing LesSchuh, results in an increased step width and a decreased degree of foot placement control, when compared to wearing regular shoes, regardless of the treadmill.

We found a significant wider step width while wearing LesSchuh, compared to wearing the regular shoe. This suggests that maintaining stability during walking is more difficult with constrained ankle moment control. As errors in foot placement control cannot be corrected for by ankle moment control anymore, increasing step width could serve as a compensation strategy (van Leeuwen et al., 2021). This is in line with our hypothesis. We also found a greater effect of LesSchuh on the split-belt treadmill compared to the C-mill. This might be explained by fear of stepping in the gap while wearing LesSchuh on the split-belt treadmill. Stepping in the gap with LesSchuh leads to a significant perturbation. To compensate, the walking strategy of the participants might have been adjusted, resulting in the greater step width. After the experiment, participants sometimes reported that they were afraid to step in the gap (note that we did not explicitly assess fear).

As expected from previous work, we also found a decreased degree of foot placement control when walking with LesSchuh, but only on the split-belt treadmill. On the C-mill, wearing LesSchuh resulted in an increased degree of foot placement control. These results are not in line with our hypothesis, because we predicted a decreased degree of foot placement control, regardless of the treadmill (Van Leeuwen et al., 2020). In the previous study, we explained the lower degree of foot placement control as indicative of how ankle moment control may not only complement foot placement by correcting errors after foot placement, but also is an intrinsic part of foot placement control itself (Zhang et al., 2020). Thus, we attributed the decrease in foot placement control to perturbed ankle moments effectively perturbing foot placement control. However, even in our previous study, we had actually hypothesized an increased degree of foot placement control when walking with LesSchuh. We reasoned that without ankle moment control to correct for foot placement errors, participants would compensate by tightening their degree of foot placement control. The latter seemed to have occurred for our current C-mill results, showing that constrained ankle moment control increases the degree of foot placement control. Clearly, the effects we previously reported were confounded by the gap between the belts. The additional task constraint of avoiding the gap, is likely to have led to the decreased foot placement control. This is in line with a lower degree of foot placement control when subjects are instructed to walk on lines projected on a treadmill (Perry & Srinivasan, 2017). Here we provide further evidence that when an additional task constraint is imposed on foot placement, the degree of foot placement control with respect to the CoM is compromised.

### 4.1. Limitations

A limitation of this study is that, apart from adjusting step width and foot placement control, other compensation strategies may have been used to maintain stability while wearing LesSchuh. One such strategy may have been increasing stride frequency, and indeed wearing LesSchuh resulted in a small but significant increased stride frequency (on average 0.004 Hz) on both treadmills (Hak et al., 2013; van Leeuwen et al., 2020b). Additionally, walking on the split-belt treadmill resulted in a significant lower stride frequency (on average 0.005 Hz) as compared to on the normal treadmill, both when wearing LesSchuh as well as when wearing the regular shoe. Perhaps, the differences in stride frequency between treadmills reflect a prioritization of different stability mechanisms on a split-belt treadmill as compared to on a normal treadmill. As foot placement is constrained by the gap, on a split-belt treadmill one may tend to prioritize stance leg control more. Ankle moment control has been shown to contribute more prominently in response to a perturbation at lower stride frequencies (Fettrow et al., 2019). This is arguable because the longer stance time at lower stride frequencies makes ankle moment control more effective. At higher frequencies, the contribution of foot placement control in response to a perturbation was higher as compared to at lower frequencies (Fettrow et al., 2019).Thus, the participants in the current study may have lowered their stride frequency on a splitbelt treadmill, to effectively prioritize ankle moment control in response to the constrained foot placement. Toeing out may have been another strategy that subjects may have used. Indeed, wearing LesSchuh resulted in significantly more toeing-out (on average 1.9°) on both treadmills (Hoogstad et al., 2022; Rebula et al., 2017). Also, walking on the split-belt treadmill resulted in a significant increase in toeing-out (on average 2.9°), wearing LesSchuh as well as wearing the regular shoe. Because of these small differences, we expect that the adjustments in toeing-out do not significantly influence our results. Lastly, despite the instructions to walk on the beam of LesSchuh, this might not have happened continuously.

### 4.2. Practical relevance & recommendations

Previous work has suggested that LesSchuh may be used as a perturbation of foot placement control, and that the aftereffects after walking with such a perturbation can be used as training (Heitkamp et al., 2019; Hoogstad et al., 2022; Reimold et al., 2020). However, the current results show that walking with LesSchuh can only be regarded as a perturbation to foot placement control if subjects walk on a split belt treadmill. During walking on a normal treadmill, foot placement control during walking with LesSchuh actually improved (which was the original hypothesis in our earlier work). It remains to be seen whether the latter adaptation maintains as an aftereffect upon returning to normal shoes and as such whether LesSchuh has a training potential on a normal treadmill. Like LesSchuh on a normal treadmill, an assistive force field improved the degree of foot placement control during application (Heitkamp et al., 2019). Yet, an aftereffect of walking in this assistive force field was a diminished degree of foot placement control compared to normal walking. Still, LesSchuh may have a training potential on a normal treadmill. Walking with LesSchuh on a normal treadmill would promote increased active foot placement control by constraining a correction mechanism for foot placement errors, rather than decreased active control due to external assistance. Therefore, if any aftereffects after walking with LesSchuh on a normal treadmill would occur, we speculate them to be more similar to those after application of a perturbing force field (i.e. a higher degree of foot placement control as an aftereffect) (Reimold et al., 2020).

For future research investigating mechanisms stabilizing gait, we recommend using a normal treadmill whenever possible. Our study showed that walking on the split-belt treadmill affected step width and foot placement control.

## 5. Conclusion

Walking with constrained ankle moment control resulted in a wider step width on both treadmills and a decreased foot placement control on the split-belt treadmill and an increased foot placement control on the normal treadmill. The increase in step width was larger on the split-belt treadmill, compared to a normal treadmill. We conclude that the effect reported in our previous study were confounded by the use of a split-belt treadmill (Van Leeuwen et al., 2020).

## 7. Acknowledgements

Sjoerd Bruijn and Moira van Leeuwen were funded by the Dutch Research Council (016.Vidi.178.014), https://www.nwo.nl/en/.

## Notes

https://doi.org/10.5281/zenodo.7188711

### Competing Interest Statement

The authors have declared no competing interest.

### Summary of Updates

Shortened the abstract and updated the link to the data repository.

